# Identification of Genes under Purifying Selection in Human Cancers

**DOI:** 10.1101/129205

**Authors:** Robert A. Mathis, Ethan S. Sokol, Piyush B. Gupta

## Abstract

There is widespread interest in finding therapeutic vulnerabilities by analyzing the somatic mutations in cancers. Most analyses have focused on identifying driver oncogenes mutated in patient tumors, but this approach is incapable of discovering genes essential for tumor growth yet not activated through mutation. We show that such genes can be systematically discovered by mining cancer sequencing data for evidence of purifying selection. We show that purifying selection reduces substitution rates in coding regions of cancer genomes, depleting up to 90% of mutations for some genes. Moreover, mutations resulting in non-conservative amino acid substitutions are under strong negative selection in tumors, whereas conservative substitutions are more tolerated. Genes under purifying selection include members of the EGFR and FGFR pathways in lung adenocarcinomas, and DNA repair pathways in melanomas. A systematic assessment of purifying selection in tumors would identify hundreds of tumor-specific enablers and thus novel targets for therapy.

## Introduction

Tumor formation is an evolutionary process driven by positive selection for somatic mutations that provide a competitive advantage to cancer cells (Nordling 1953; Nowell 1976; Greaves and Maley 2012). While positive selection drives phenotypic change, it only enriches for a miniscule fraction of the mutations in tumor genomes (Lawrence et al. 2013; Lawrence et al.2014). During species evolution, most newly arising mutations are deleterious, and are eliminated by negative (or purifying) selection before they can become substitutions fixed in the population of individuals (Kimura and Ohta 1974; Kimura 1991; Zollner et al. 2004; Kiezun et al. 2013). In principle, negative selection could also impact cancer evolution (McFarland et al. 2013; McFarland et al. 2014), and there is evidence of purifying selection in hemizygous regions of cancer genomes (Van den Eynden et al. 2016). However, the extent to which this force shapes the pattern of somatic mutations in tumors is not known. In this study, we provide evidence that purifying selection is widespread in cancer genomes and acts to remove mutations from genes that contribute to the survival or growth of cancer cells. In this way, the pattern of mutations in patient tumors reveals the vulnerabilities of human cancers *in vivo*.

## Results

### Genes that are expressed or essential have fewer missense mutations (substitutions)

If purifying selection were significant during tumor evolution, it would reduce overall substitution rates by preventing the fixation of deleterious somatic mutations in genes contributing to tumor growth. To examine this possibility, we analyzed the mutational profiles of 5057 tumors of diverse cancer types sequenced by The Cancer Genome Atlas (TCGA) (Weinstein et al. 2013). Since genes can only impact tumor growth if they are expressed, our first analysis was to compare substitution rates between expressed and non-expressed genes (Figure 1A). Each gene’s exon mutation rate was normalized relative to its intron mutation rate; this controlled for gene-to-gene variations in mutation rates arising from differences in chromatin accessibility and early-vs-late replication times, among other position factors (Lawrence et al. 2013) (Figure 1B). After controlling for all of these effects, expressed genes had significantly fewer substitutions than non-expressed genes across three tumor types— with a 57% reduction in melanomas (p<10^−20^), a 51% reduction in lung adenocarcinomas (p<10^−20^), and a 14% reduction in colorectal adenocarcinomas (p<10^−20^) (Figure 1A). Absent this reduction, we estimate there would have been 167–416 additional mutations in the exons of expressed genes per tumor, depending on the cancer type. This depletion of missense mutations is similar to the 66-83% of missense mutations observed to impact a protein’s functionality, based on experimental mutagenesis (Rockah-Shmuel et al. 2015).

Transcription-coupled repair (TCR) (Hanawalt and Spivak 2008) has been previously reported as a mechanism through which mutations are eliminated from expressed genes. To quantify TCR’s effects, we compared substitution rates between transcribed (template) and non-transcribed (coding) strands in melanomas and lung adenocarcinomas. As expected, TCR lowered overall substitution rates in expressed genes. However, there was a 31-45% additional reduction that could not be accounted for by TCR (Supplemental Figure S1). These findings were consistent with a model in which mutations were being eliminated by purifying selection prior to their fixation.

**Figure 1.**
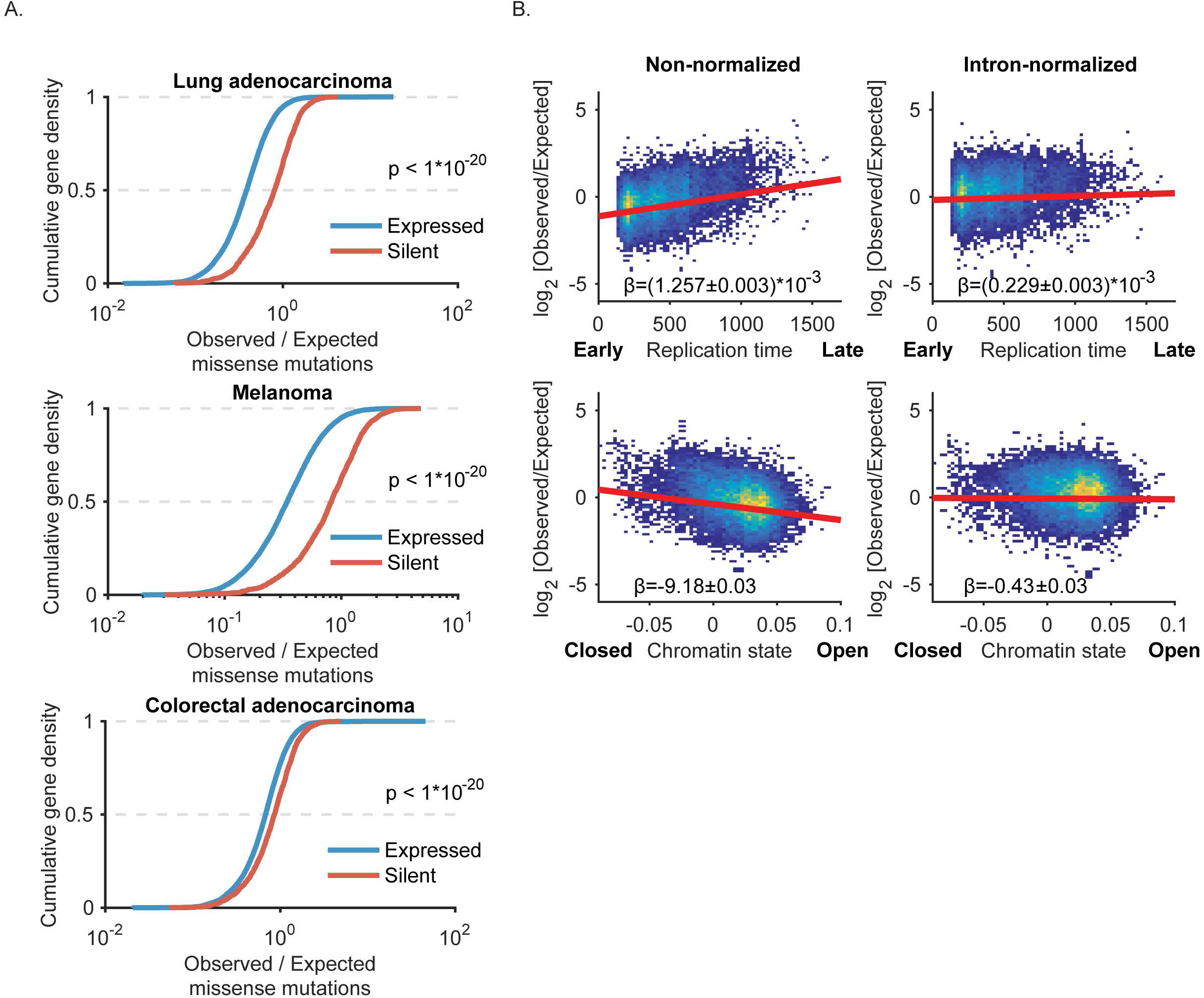
Genes that are expressed have fewer missense mutations (substitutions). (a) Cumulative distribution of observed missense mutations/ expected (intron normalized) in expressed vs. silent genes in melanomas (n = 290), lung adenocarcinomas (n = 533) and colorectal adenocarcinomas (n = 489). For each tumor type, expressed genes were defined as having an estimated transcript count > 8 in95% of tumors. Statistical significance was assessed using the Wilcoxon rank-sum test. **(b)** Expected mutation rates from intron mutations controls for various mutation covariates in lungadenocarcinomas, including %GC, replication timing, and chromatin accessibility. A linear regression is plotted of logobserved over expected mutations for non-normalized and intron-mutation rate normalized based expected. The slope ofthe regression ( ) is displayed with the 95% confidence interval. Color corresponds to density of genes in the scatter plots.

### Amino acid substitutions with similar physicochemical traits are more acceptable during both tumor microevolution and species macroevolution

Mutations resulting in the substitution of amino acids with similar physicochemical properties (conservative substitutions) are less likely to be deleterious to protein function, relative to non-conservative substitutions (Grantham 1974; Kimura and Ohta 1974). If this were the case in tumors, purifying selection should act less strongly on mutations resulting in conservative amino acid substitutions. To test this prediction, we segregated mutations into classes based on the amino acid substitutions that they generated. In total, there were mutations in all of the 150 substitution classes that are possible by mutating a single base pair in codons. We quantified the strength of negative selection on each mutation-substitution class to identify pairs of amino acids (A1, A2) that were most readily substituted in either direction (A1 -> A2 and A2 -> A1) in tumors (Figure 2A,B, Supplemental Table S1). This analysis identified several subsets of amino acids with similar physicochemical properties that were interchangeable in tumors: the hydrophobic amino acids isoleucine, leucine, valine, and methionine; the positively charged amino acids arginine, histidine, and lysine; and the positively charged and positive-polar amino acids arginine and glutamine. The analysis also identified several amino acids with similar structures but differing charges that were interchangeable (Gln⇔Glu and Asp⇔Asn), suggesting that such substitutions might minimize steric hindrances and be frequently tolerated. We conclude that mutations resulting in conservative substitutions were less often eliminated by purifying selection in tumors— presumably because they were less likely to disrupt protein folding or function.

**Figure 2.**
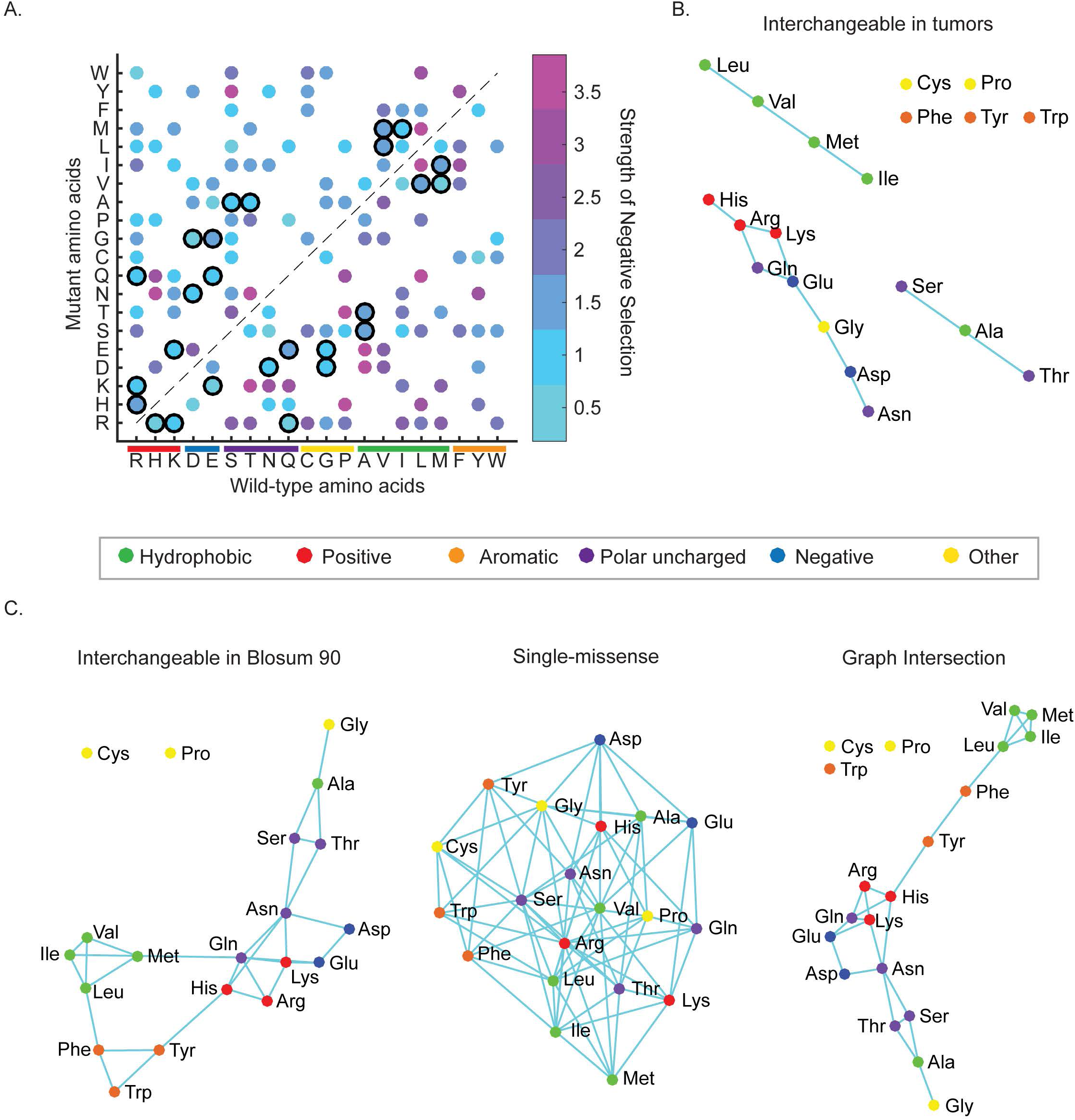
Amino acid substitutions with similar physicochemical traits are more acceptable during both tumor micro-evolution and species macroevolution. (a) Heat map showing the strength of negative selection on each observed pairwise amino acid substitution. Bold outlines highlight substitutions between interchangeable amino acids. Negative selection strength was quantified as described in methods. **(b)** Graph depicting amino acids interchangeable during tumor microevolution,color-coded based on their chemical properties. **(c)** Graph depicting amino acids interchangeable during species macroevolution, defined as substitutions with a BLOSUM90 score0 (left). Graph depicting all amino acid substitutions that are possible with a single missense mutation (center). The graph corresponding to the intersection of these two graphs is also shown (right). The intersection between conservative transitions in tumors and in BLOSUM, has a p-value of 7.64*10^−6^, as determined from the hypergeometric distribution.

The constraints imposed on protein folding and function during tumor microevolution might in principle be comparable to those imposed during the macroevolution of species. We therefore compared the amino acid substitutions that were tolerated in tumors with those that were most commonly tolerated across macro-evolutionary time scales. Surprisingly, we found that interchangeable amino acids identified using BLOcks of Amino Acid SUbstitution Matrix (BLOSUM; (Henikoff and Henikoff 1992)) analysis— which quantifies substitutions within highly conserved protein domains across millions of years of species evolution— were nearly identical to those identified in the tumor analysis (p < 7*10^−6^) (Figure 2C). However, this concordance was only observed if: (1) the macro-evolutionary analysis was performed for closely related proteins (BLOSUM90, but not BLOSUM45/62), and (2) the BLOSUM90 amino acid substitutions were limited to those that are possible by mutating a single DNA base in codons; both of these constraints reflect the fact that the substitution rates in tumors are much lower than those observed in comparisons across species. Moreover, this analysis revealed that several substitutions that were well tolerated in tumors, which could not be understood on the basis of their physicochemical traits (e.g. Glu ⇔Lys, Ser ⇔ Ala,Alar ⇔Thr), were also more tolerated across the macro-evolutionary time scales associated with speciation, suggesting that they are in fact more permissible than others (Figure 2B, C). After considering these findings together with the functional observations above, we concluded that purifying selection has a significant role in shaping the global constellation of substitutions (fixed mutations) found in tumors.

### Purifying selection targets genes that are important for tumor growth

Since these findings established that negative selection occurred at a genome-wide scale in tumors, we next asked whether we could identify individual genes that were substrates of purifying selection. We found genes associated with essential processes, such as transcription (*MED15*, *MED19*) and cell division (*ANAPC2*, *CEP72*), Supplemental Table S2) to be under purifying selection in tumors. However, we could not detect evidence of purifying selection in genes with too few mutations across the sequenced tumors. To work around this, we looked for purifying selection in sets of genes with known biological functions (Liberzon et al. 2011). We found that genes which function in essential cellular processes— e.g. RNA metabolism and DNA replication— are under the strongest purifying selection across all tumor types (Figure 3A, B; Supplemental Table S3). In addition to these gene sets showing a depletion of mutations, we found that in each set under purifying selection, the majority of genes showed fewer mutations than expected (Figure 3C).

To support our observation that genes under purifying selection were enriched in essential cellular processes, we examined if these genes were known to be essential when perturbedAssembling the results of three pooled CRISPR screens (Hart et al. 2015; Wang et al. 2015; Tzelepis et al. 2016), we found that genes under purifying selection are more often essential in most tested cell lines, compared to genes not under selection (Figure 3D). We also found that purifying selection has a strong power to find essential genes (Figure 3E). This showed that genes under purifying selection in tumors are functionally essential, suggesting it reveals genes important for tumor growth or survival.

Although many genes under purifying selection across tumors are essential, such genes are not likely to be good targets for treating cancer, as they likely also have essential functions in normal cells. To get around this, we aimed to identify genes under increased selection in particular tumor types, relative to other tumors. To identify genes under increased purifying selection in specific tumor types, we developed a statistical approach that controlled for differences in the stage of tumor evolution, gene-specific variations in mutation rate within tumors, and genome-wide variations in mutation rates across tumors. Using this method, we could identify genes under purifying selection in specific tumor types— e.g. in lung tumors versus all other tumor types.

**Figure 3.**
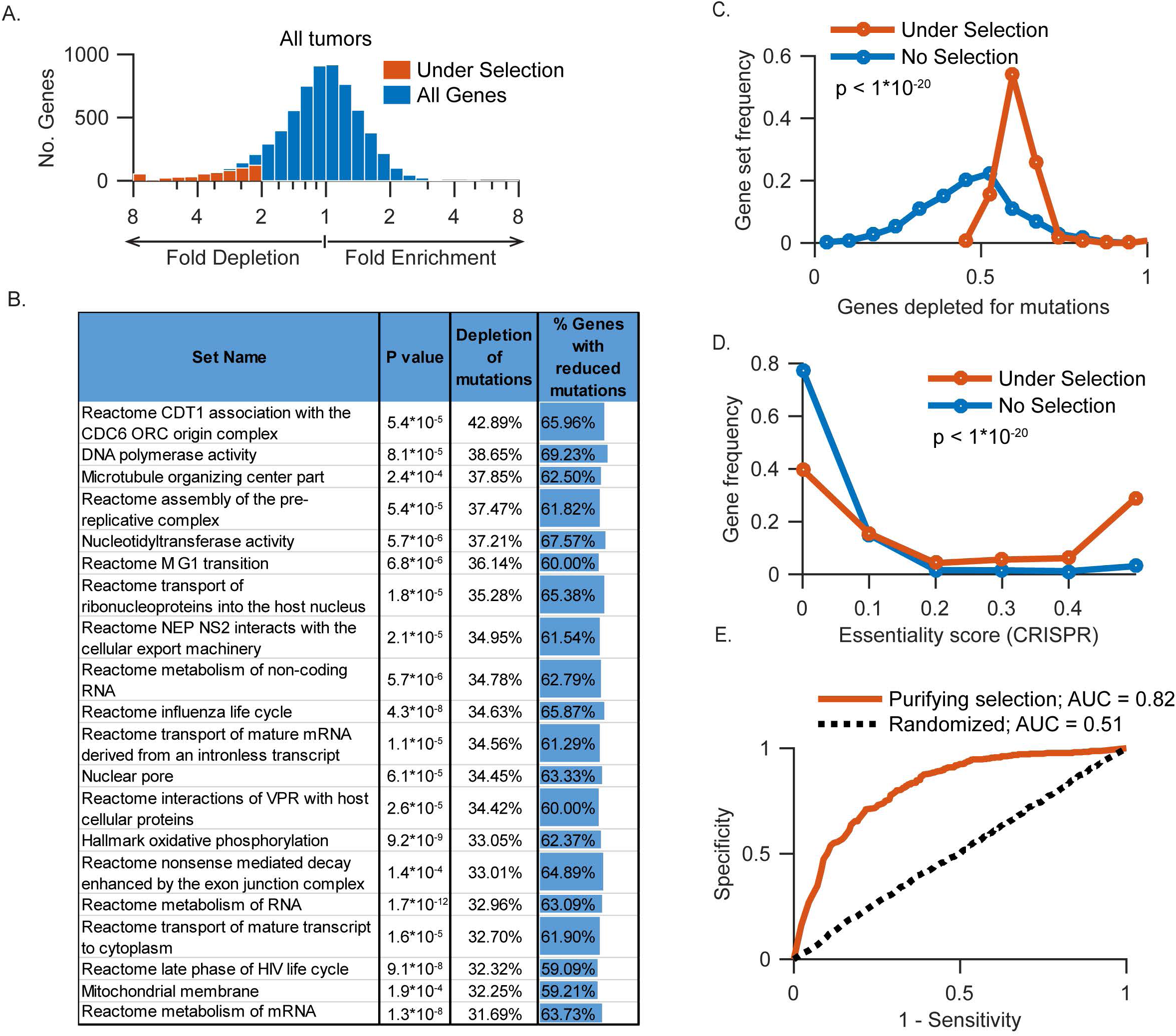
Purifying selection targets genes that are important for tumor growth. (a) Histogram showing the number of genes depleted for substitutions across all tumors (vs expectation from intron mutations). Genes with significant depletion (p<0.01 and >2-fold) are shown in red, and all genes in blue; all genes shown have:::10 expected mutations and are expressed in all tumors. **(b)** Shown are the top 20 significant (Q<0.01) gene sets ranked by the depletion of mutations vs expected. P values were determined through sampling (see Methods). “% Genes with reduced mutations” represents the proportion of genes within each set with fewer mutations than expected. **(c)** Most genes in gene sets under purifying selection have fewer mutations than expected. Statistical significance was assessed using the Wilcoxon rank-sum test. **(d)** Many genes under purifying selection across tumors are essential. Genes are binned by the proportion of cell lines in which they were deemed essential based on 3 published pooled CRISPR screens (see methods). Statistical significance was assessed using the Wilcoxon rank-sum test. **(e)** Receiver-operator characteristic (ROC) curves showing the preditive value of purifying selection, based on genes’ estimated p values, for identifying essential genes. Also shown are random genes. Essential genes were identified in a published pooled CRISPR screen in cancer cell lines. AUC, area under the curve.

In lung adenocarcinomas, we identified 508 genes as strong substrates for purifying selection, enriched in 11 of the pathways in the network data exchange database (NDEx) (Supplemental Figure S2, Supplemental Table S4, Supplemental Table S5) (Pratt et al. 2015). These included: 11 genes in pathways related to EGFR signaling (ERBB2/ERBB3, EGFR internalization, ERBB1 receptor proximal pathway, p<3x10^−3^, Supplemental Figure S2), and the AXL kinase. These pathways are both targeted by approved therapies for lung cancers: erlotinib/gefitinib (EGFR) and crizotinib (MET/AXL). Our analysis also identified *FGFR3*, a key driver of non-small cell lung cancer (NSCLC), which is activated by mutation in 6-8% of NSCLCs and is currently being explored as a therapy target (Semrad and Mack 2012; Liao et al. 2013; Yin et al. 2013; Wang et al. 2014; Tiseo et al. 2015).

In cutaneous melanomas, we identified 848 genes that were targets of purifying selection. Consistent with the established role of UV-induced damage in this cancer type, these included 27 genes in key pyrimidine dimer repair pathways: nucleotide-excision (*ERCC2*, *ERCC5*), base excision (*APEX2, POLE*), mismatch repair (*RFC1, RFC4*), and trans-lesion replication (*REV1, REV3L*) (Figure 4A,C, Supplemental Table S6, Supplemental Table S7). While UV does not directly cause double-stranded breaks (DSBs), such breaks arise indirectly during NER and are the primary cause of cell death (Wakasugi et al. 2014). Consistent with this, we identified a number of genes that repair DSBs in the ATM and Fanconi Anemia pathways (ATM & FANCONI pathways, p <4x10-5; Table S7)— including two members of the core Fanconi Anemia complex (*FANCC, FANCL*), *ATM* and its phosphorylation target CHK2, and 2/3 proteins in the MRN complex (NBN and RAD50) (Figure 4C). We also identified all four components of the cohesin complex (SMC1A, SMC3, STAG2, RAD21), which, independently of its role in mediating sister chromatid cohesion, is phosphorylated by ATM and required for repairing DNA DSBs by homologous recombination (Kim et al. 2002; Yazdi et al. 2002; Kong et al. 2014). Collectively, these findings indicate that purifying selection preserves the function of DNA repair pathways in melanomas. Because many of these pathways have established roles in promoting resistance to the DNA damage caused by radiation and chemotherapies (Reed 1998; Helleday 2010; Begg et al. 2011; Pennington et al.2014; Dai et al. 2015); this might explain why such therapies are almost completely ineffective when applied to melanomas.

**Figure 4.**
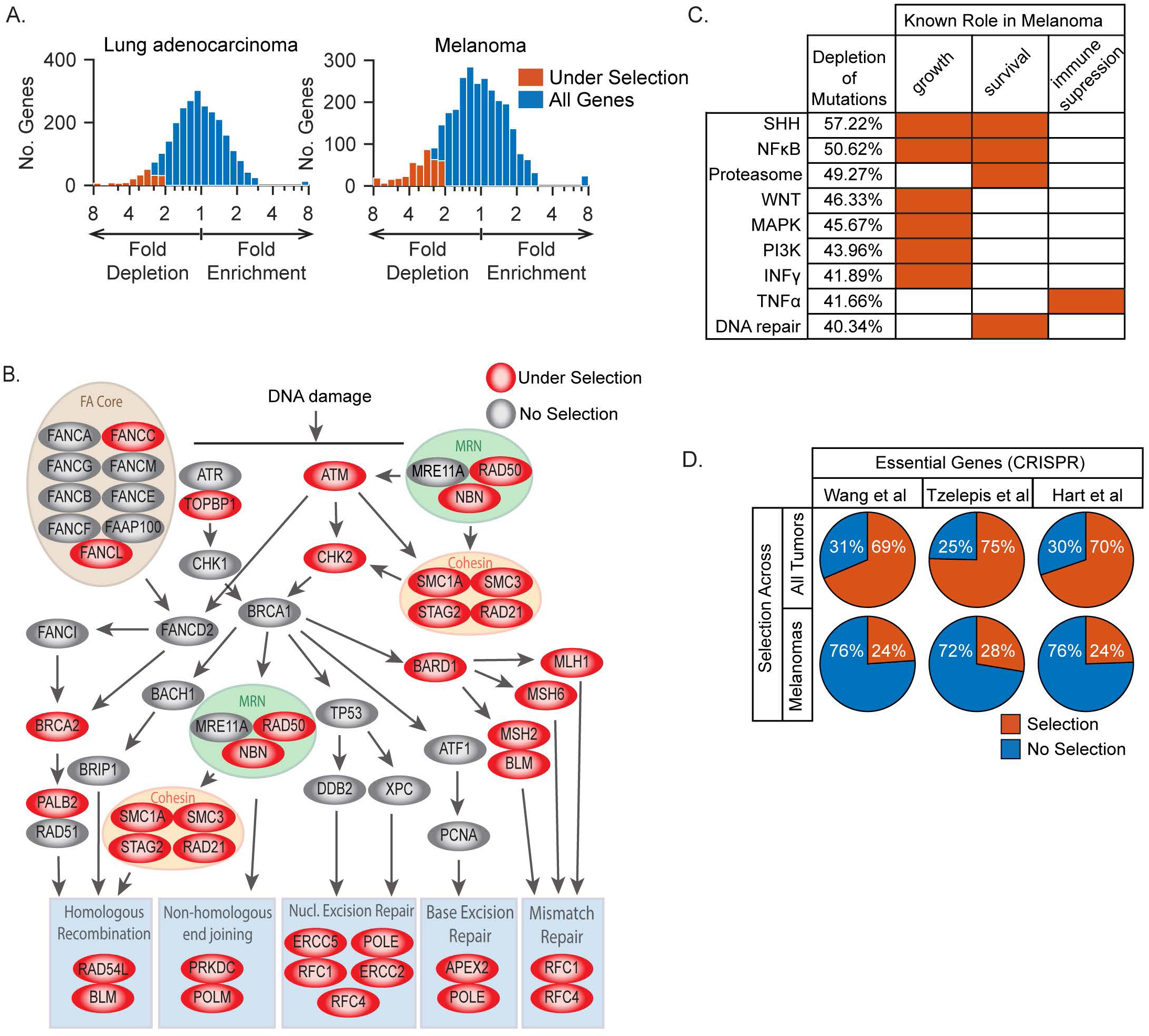
Purifying selection reveals tumor type-specific vulnerabilities. (a) Histograms showing the number of genes depleted for substitutions across tumors of the indicated type relative to other tumor types. Genes with significant depletions (p<0.02 and >2-fold) are shown in red, and all genes in blue; all genes shown have at least 10 expected mutations and are expressed in all tumors. **(b)** Shown are DNA repair pathways, highlighting genes (red) that are targets of increased purifying selection in melanomas (p < 0.02 with >2-fold depletion). **(c)** Pathways under more purifying selection in melanomas are known to be important for growth, survival, immune supression, or metastasis in melanomas. Shown are pathways identified from gene sets with increased purifying selection in melanomas relative to othertumor types (Q < 0.05 and > 50% depletion of mutations; pathways shown represent 519 / 882 genes under selection). “Depletion of mutations” represents the average percent depletion of mutations in genes in sets corresponding to each pathway. The depletion in melanomas is calculated relative to the expected number of mutations, based on the number of mutations observed in other tumor types. **(d)** Fewer essential genes are under increased purifying selection in melanomas, compared to genes under purifying selection across all tumor types. Each pie chart shows the proportion of essential genes under purifying selection in all tumor types, or under increased selection in melanomas. Genes essential in cancer cell lines were identified from published pooled CRISPR screens.

We were again able to use identify purifying selection on sets of genes with known biological function, this time looking for increased selection in a particular tumor type. Gene sets under increased purifying selection in melanomas are related to a number of pathways active in processes known to be important in melanomas (Figure 4B, Supplemental Table S8). Describing 599/927 genes in these sets, we found many known pathways involved in melanoma growth and survival, such as the sonic hedgehog, WNT, NF?B, PI3K, EGFR, and INF? pathways, and the proteasome (Rubinfeld et al. 1997; Ueda and Richmond 2006; Boone et al. 2011; Yaguchi et al. 2012; Jalili et al. 2013; Selimovic et al. 2013; Webster and Weeraratna 2013; Gross et al. 2015). We also found a pathway required for immune suppression in melanomas, TNFα (Wang et al. 2016).

Importantly, genes under increased purifying selection in melanomas are less likely to be generally essential for cell viability when compared to genes under purifying selection in all tumors (Figure 4D).

When attempting to extend this analysis to other tumor types, we found that there were not enough passenger mutations identified to provide the statistical power needed for the analysis. How much more benefit would be obtained by sequencing additional tumors? Using numerical simulations, we estimated the number of new genes that would be discovered by sequencing 500 to 3000 additional tumors of each cancer type (Supplemental Figure S3). For all tumor types, sequencing no more than 500-3000 additional tumors would be sufficient to discover nearly all of the genes under purifying selection that have yet to be identified. In addition, we established the optimal combination of tumor types to sequence that would maximize the number of new genes discovered as substrates of purifying selection (Supplemental Figure S3).

## Discussion

These findings show that purifying selection significantly influences the pattern of mutations in cancer genomes, reducing the rate at which substitutions accumulate in genes that are important for tumor growth. We propose calling genes under purifying selection in tumors’enablers’, to distinguish them from recurrently mutated ’drivers’ — i.e., tumor-suppressors and oncogenes. Our findings indicate that many enablers are tumor type-specific, and are therefore not likely to be generally required for the survival of all cell types; however, it may also be that there are tissue-specific differences in essential genes. Enablers that are tumor type-specific could arise through cell type-specific requirements or through synthetic interactions with genetic and metabolic alterations associated with tumor growth, as recently reported (Kryukov et al. 2016; Mavrakis et al. 2016). Using signatures of purifying selection to discover enablers provides an exciting opportunity to systematically identify hundreds of new vulnerabilities of cancer. As the vulnerabilities of human tumors will remain opaque to direct experimentation,and only approached by models, our observation of purifying selection in cancers allows an unprecedented view into the dependencies of human cancers *in vivo*.

## Methods

### Data Availability

All post-analysis data are included in this manuscript in supplemental tables. All data analyzed were obtained from other sources as follows.

### Tumor mutation data

Mutation Annotation Format files for 11 tumor types generated by The Cancer Genome Atlas (TCGA) were downloaded from the Broad Firehose (Broad Institute TCGA Genome Data Analysis Center (2015): Firehose stdata _2015_11_01 run. Broad Institute of MIT and Harvard. doi:10.7908/C1571BB1). Tumor types downloaded were lung adenocarcinoma (533 tumors), cutaneous melanoma (290 tumors), colorectal adenocarcinoma (489 tumors), bladder urothelial carcinoma (395 tumors), breast invasive carcinoma (977 tumors), glioma (796 tumors), uterine corpus endometrial carcinoma (248 tumors), head and neck squamous cell carcinoma (510 tumors), liver hepatocellular carcinoma (198 tumors), prostate adenocarcinoma (332 tumors), and stomach adenocarcinoma (289 tumors). Mutations were filtered to remove all but single base-pair missense mutations in exons.

Non-coding (intron) mutation data from were acquired from published analyses. (Lawrence et al. 2013)

### RNA data

Level 3 normalized RNA sequencing data quantified with RNA-Seq by Expectation Maximization (RSEM)(Li and Dewey 2011) were downloaded from the Broad Firehose (Broad Institute TCGA Genome Data Analysis Center (2015): Firehose stdata 2015_11_01 run. Broad Institute of MIT and Harvard. doi:10.7908/C1571BB1). These data are quartile-normalized RSEM count estimates.

### Gene-length and sequence information

Gene length information was downloaded from UniProt (http://www.uniprot.org/), and coding sequences were downloaded from BioMart (http://www.biomart.org/).

## Calculations

### Mutation rates in expressed and non-expressed genes

For tumor *t* ∈ tumor type *T*_i_∈ *T,* where *T* = {Lung adenocarcinoma, skin cutaneous melanoma, colorectal adenocarcinoma} (see Tumor mutation data, above); and for gene *g* ∈ *G,* where *G* = all sequenced genes; and where *Lg* = the length of gene *g* in amino acids (a.a.s);

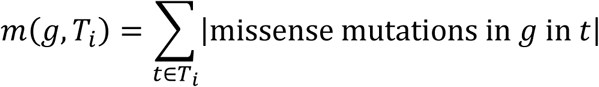

Where

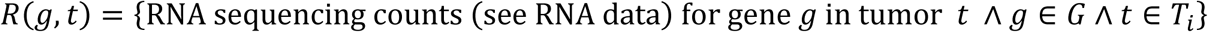

Define expressed genes

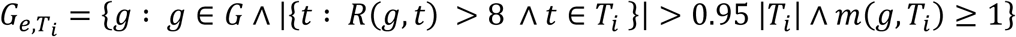

and not-expressed genes as

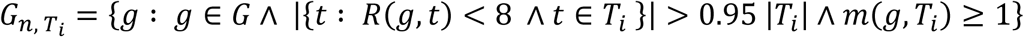

Determine an expected number of mutations for each gene by means of the gene’s relative non-coding mutation rate, the average mutational rate in not expressed genes, and the length of the gene’s coding sequence:

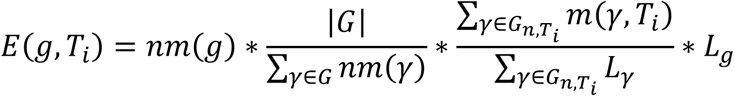

Where *nm(g)* = the non-coding mutation rate for gene *g* calculated from published whole-genome sequencing of tumor samples (Lawrence et al. 2013).

To calculate the significance of the depletion in mutations in expressed genes,

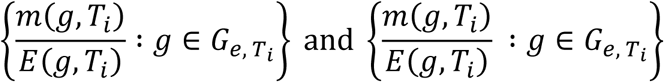

were compared with a two-tailed Wilcoxon Rank-Sum test.

The proportion of mutations depleted in expressed genes relative to not expressed genes was calculated as

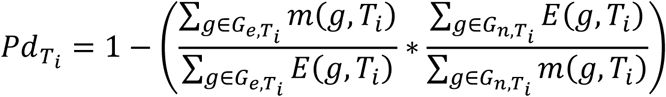

The number of additional expressed mutations expected in sequenced tumors was calculated as

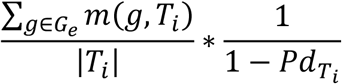

### Determining the effect of intron mutation rate controls on mutation rate covariates

Using the intron mutation rate to estimate the background mutation rates of genes should ideally control for known gene mutation rate covariates, including replication time, chromatin accessibility, and GC nucleotide percentage. Where *Ti*= lung adenocarcinomas, observed and expected (intron-normalized) missense mutations were calculated for each expressed gene as in “Mutation rates in expressed and non-expressed genes,” above. Replication time and chromatin accessibility of each gene were accessed from a published source (Lawrence et al. 2013). The %GC nucleotides of each gene was determined from each gene’s coding sequence. To determine these relationships before controlling via the intron mutation rate, an expected number of mutations was calculated for each gene assuming a uniform mutation rate, or *E*^0^(g):

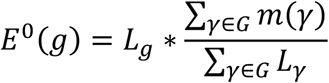

For both the intron-normalized expected and the uniform mutation rate expected, Each covariate score for expressed genes was plotted against the log2 observed / expected mutations of those genes, and a linear regression determined.

### Estimating the effects of transcription-coupled repair

To estimate the effect of transcription-coupled repair, mutation rates were quantified in the transcribed and not-transcribed strands. For each missense mutation µ, define the starting base 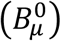 and ending base 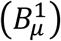, and its indistinguishable complement with starting base 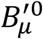 and 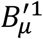. There are six kinds of recognizable base-pair transitions, as some are indistinguishable from a mutation in the opposite strand.

For G>T mutations in lung adenocarcinomas and C>T mutations in melanomas, mutation rates were calculated on a gene-by-gene basis in expressed and not-expressed genes.

Define 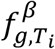 as the number of mutations in of the class *β*^0^ > *β*^1^e.g. C>T) in the transcribed(template) DNA strand of gene *g* in tumor type *Ti*, and 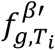 ' as the mutations in the class *β*^0^ > *β*^1^in the not-transcribed (coding) DNA strand of gene *g* in tumor type *Ti*:

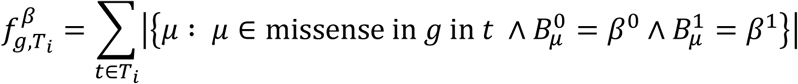

and

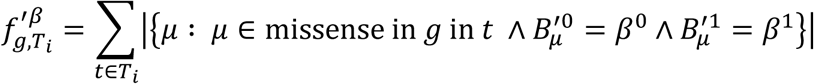

Also define 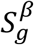 and 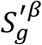 as the number of sites that could mutate in the transcribed (template)DNA strand and not-transcribed (coding) DNA strand of gene *g* respectively, or

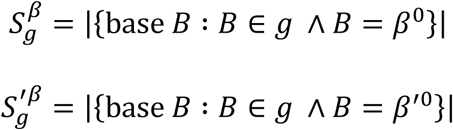

Determine an expected number of mutations for each gene, and for each strand, by means of the gene’s relative non-coding mutation rate, the average mutational rate in expressed genes, and the length of the gene’s coding sequence of the base in question (for the template strand)or its complement (for the coding strand):

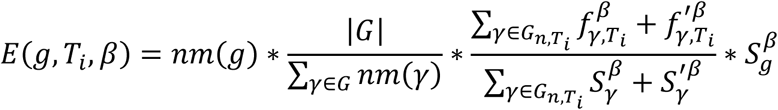

And:

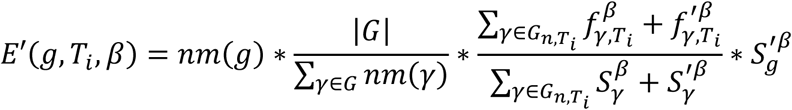

To test the relative mutation rates of the non-transcribed (coding) strand,

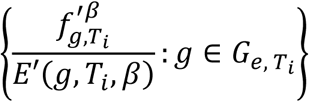

and

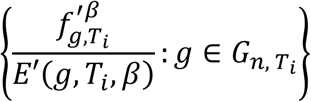

were compared with a two-tailed Wilcoxon Rank-Sum test.

The percent depletion of mutations in expressed genes remaining after controlling for transcription coupled repair and noncoding mutation rates was computed by comparing the mutation rate of the not-transcribed strand of expressed genes and the mutation rate of the not-transcribed strand of not expressed genes, or:

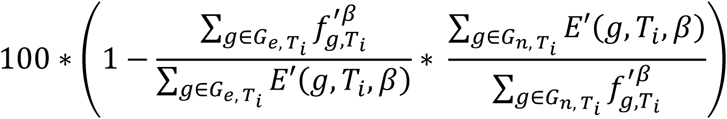

For each observable transition β (e.g. G>A), where the starting base is defined as β^0^ (or its complement *β*^0^), and the ending base as β^0^ or its complement *β*^0^, the percent depletion of mutations in expressed genes was calculated in each strand. For the not-transcribed (coding)strand, this rate in tumor type *T*_*i*_ is:

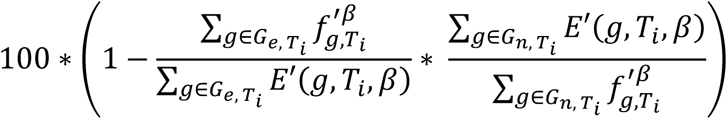

While for the transcribed (template) strand, this rate in tumor type *Ti* is:

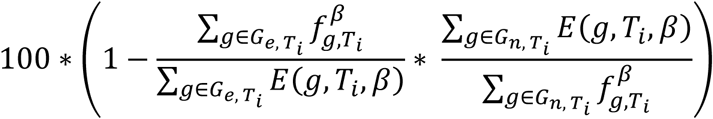

### Finding conservative amino acid transitions from cancer mutation data

To determine the strength of selection on individual amino acid (a.a.) substitutions, a.a. substitution rates in lung adenocarcinomas from TCGA were examined. *T* is defined as the set of sequenced lung adenocarcinomas (see Tumor mutation data, above), and *G* is the set of sequenced genes.

Where

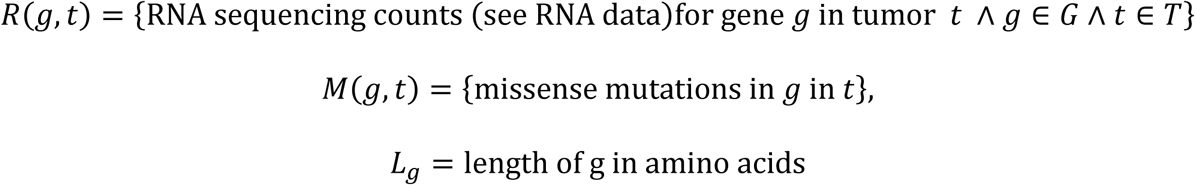

define expressed genes as:

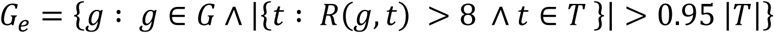

and not expressed genes as

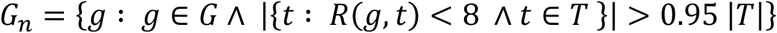

Call

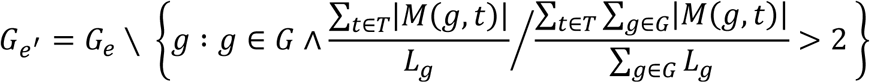

Determine the matrix of transitions between each a.a. in expressed genes

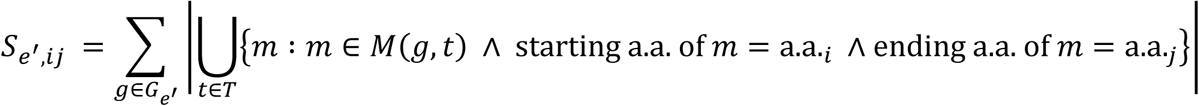

and the matrix of transitions between each a.a. in not expressed genes

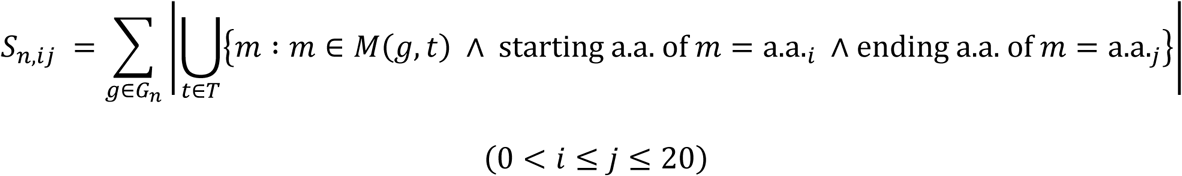

Where

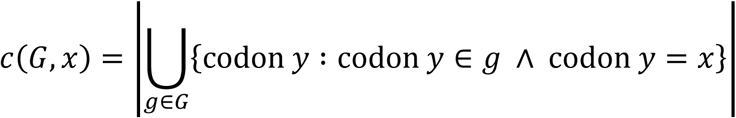

And

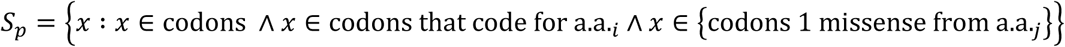

Compute the matrix of starting codon counts in expressed genes:

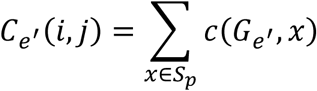

and compute the matrix of starting codon counts in not-expressed genes:

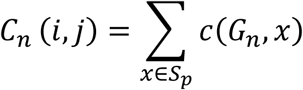

for (0<*i* ≤ *j* ≤ 20),

giving *S*_e_′, *S*_*n*_, *C*_*n*_′ and *C*_*n*_ a size of 20 x 20.

Compute the average depletion of substitutions γ such that

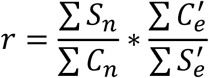

Use *r* to calculate an expected rate for each amino acid substitution in expressed genes, or

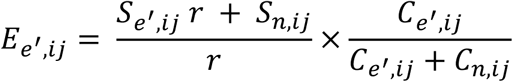

and in not-expressed genes

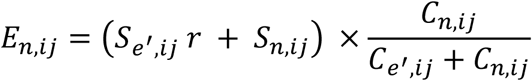

Using the expected and observed matrices, calculate a ?^2^ statistic for each substitution, stored as matrix *X* such that

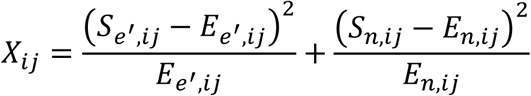

Use *X* to compute p values with the *χ*^2^ test with one degree of freedom, giving matrix *P*, where

*Pi,j* = the p value calculated from *Xij*

Also calculate matrix *F* where

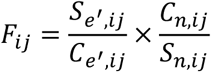

a.a.i and a.a.j are called substitutable if

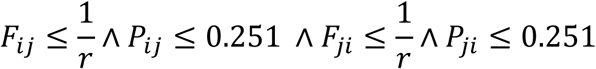

### Finding conservative amino acid transitions from BLOSUM

Substitutable amino acids from BLOSUM 90 were identified as pairs of amino acids with BLOSUM log-odds scores > 0. The significance of the overlap between substitutable amino acids identified from BLOSUM and those identified in tumors was calculated with the CDF of the hypergeometric distribution.

### Identifying genes under purifying selection in multiple tumor types

To find genes under purifying selection in multiple tumor types, data from melanomas, lung adenocarcinomas, colorectal adenocarcinomas, liver hepatocellular carcinomas, gliomas, and breast invasive carcinomas were used (forming set *T*). First, genes were only included in the analysis if they were called expressed in all tumor types, where

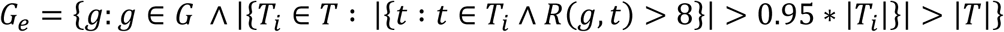

An expected number of mutations was computed for each gene, or *E*(*g*), based on eacgene’s non-coding / intron mutation rate in tumors subjected to whole-genome sequencing(Lawrence et al. 2013):

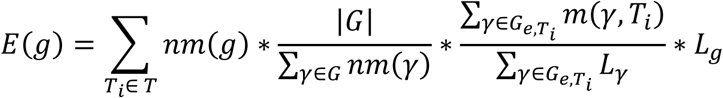

Where *Lg* = the length of gene *g* in amino acids, *nm(g)* = the non-coding mutation rate for gene *g* calculated from published whole-genome sequencing of tumor samples (Lawrence et al. 2013), and

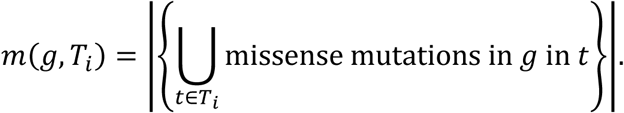

In this way, recurrent mutations (the same missense mutation observed more than once) within each tumor type were dropped from the analysis.

The expected number of mutations was compared to observed number of mutations, where

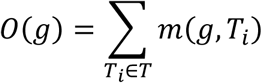

Genes were identified as under purifying selection (*N*) in these tumor types if they passed a p value and fold change cutoff:

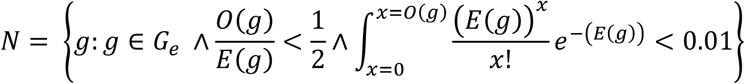

### Identifying gene sets under purifying selection in multiple tumor types

Gene sets were obtained from the Molecular Signature Database (Subramanian et al. 2005) version 5.1; sets examined were from the hallmark, canonical pathways, BioCarta, KEGG, Reactome, and GO subsets of the Molecular Signature Database, totaling 2834 sets, making 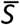 with set *S* ∈ 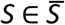. To find sets under purifying selection, mutations in these sets were examined in the melanoma, lung adenocarcinoma, and colorectal adenocarcinoma tumor types {*Ti* ∈ *T*}. For each tumor type, expressed genes were defined as genes

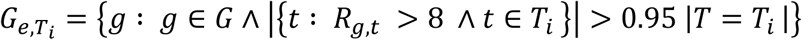

Sets were filtered so that they only contained genes with mutations in these tumors, so

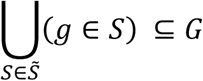

and so that

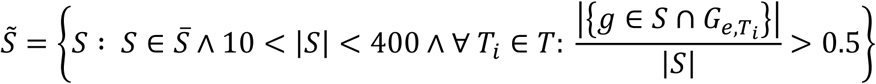

An expected number of mutations was computed for each set, where

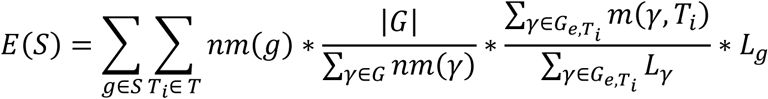

Where *Lg* = the length of gene *g* in amino acids, *nm(g)* = the non-coding mutation rate for gene *g* calculated from published whole-genome sequencing of tumor samples (Lawrence et al. 2013), and

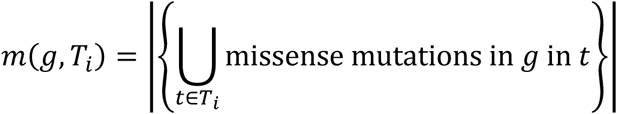

An observed number of mutations was also computed for each set, or *O*(*S*), where

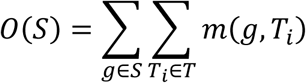

The difference between the observed and expected numbers of mutations for each set was determined through the CDF of the Poisson distribution, where

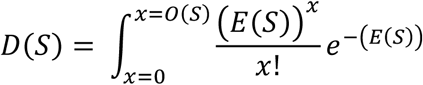

To determine the significance of the depletion of mutations for each set, 1*104 random sets 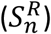 were generated for each set size, drawing from those genes in the union of all gene sets, so that

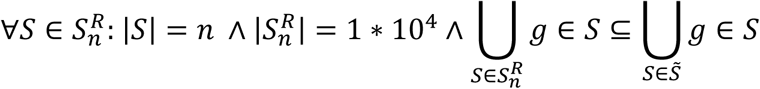

The significance of each set’s depletion in mutation was evaluated by computing a p value, or *p*(*S*), based on the depletion of random sets of the same size, so that.

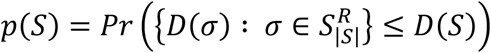

As many sets were depleted beyond even *104* random sets, an estimated p value was computed by regressing the randomly sampled sets. For each size set, the –log10 quantiles of those random sets with *p(S)* < 0.01 were fit with a linear regression vs the –log10(percentiles) that at which the quantiles were evaluated. This regression, generating for each set size slope *b* and constant *c*, was used to compute the revised p values, *P*(*S*), where

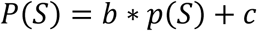

To correct for multiple hypothesis testing, a q-value was calculated using the method of Benjamini and Hochberg. (Benjamini and Hochberg 1995)

### Essentiality analysis of genes under purifying selection

The impact on cancer cell line growth of CRISPR-mediated knockout has been previously published (Hart et al. 2015; Wang et al. 2015; Tzelepis et al. 2016). In each of these three published pooled CRISPR screens, the investigators used differing methods to call whether a gene was essential. In each screen, a gene was called essential or not essential in each tested cell line. From this data, for each gene *g*, a score *C*(*g*) was recorded, or the proportion of tested cell lines in which this gene was deemed essential, based on the published results, from these three screens.

Gene sets under purifying selection across all tumor types were identified as above (“Identifying gene sets under purifying selection in all tumor types”). Sets under purifying selection (*Sp*) weredefined to be sets with q-values < 0.05, and an observed / expected number of mutations <0.8. Genes under purifying selection (*Gp*) were defined as the union of genes in sets under purifying selection (*Sp*), or *G*_*p*_ = *U*_sεs__p_ 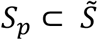 where 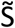 represents those sets examined from the

Molecular Signature Database (see above). Genes under purifying selection (*Gp*) was then compared to genes not under purifying selection (*Gnp*), where *G*_*np*_ = *U*_sεs__np_ *g* ε *S* 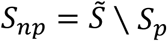

The essentiality of genes under purifying selection (*Gp*) was compared to the essentiality of genes not under purifying selection (*Gnp*); the essentiality of each gene was defined based on its score *C*(*g*), as defined above, representing the proportion of tested cell lines in which this gene was deemed essential. To calculate the significance of the difference in essentiality between these groups of genes,

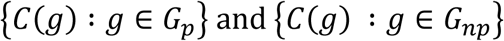

were compared with a two-tailed Wilcoxon Rank-Sum test.

To determine the utility of purifying selection for finding essential genes, a receiver-operator characteristic curve was generated using genes ranked by their revised P values (see above). Each gene was given the lowest revised P value of the gene sets examined in which it was part. Genes that were called true positives (essential) were defined as genes that were deemed essential in > 5 / 7 examined cell lines in a pooled CRIPSR screen(Tzelepis et al. 2016).

### Identifying genes under tumor type-specific purifying selection

First, genes were only included in this analysis if they were not called unexpressed in all tumor types, where

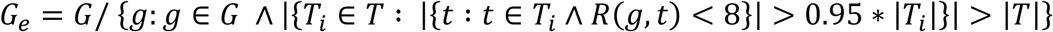

Genes with an increased mutation rate across tumors were also filtered out. An expected number of mutations was computed for each gene, or *En*(*g*), based on each gene’s noncoding/ intron mutation rate in tumors subjected to whole-genome sequencing(Lawrence et al. 2013):

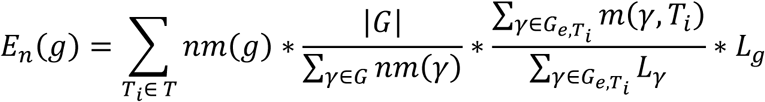

Where *Lg* = the length of gene *g* in amino acids, *nm(g)* = the non-coding mutation rate for gene *g* calculated from published whole-genome sequencing of tumor samples (Lawrence et al. 2013), and

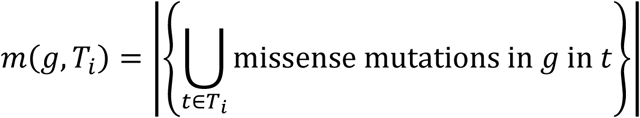

To identify genes under negative selection in a particular tumor type relative to other tumors, a different expected value was computed based on the mutation rate in all other tumors. For tumor type *Ti* ∈ *T* where *T* = {*T*1, *T*2, … *T*11} or all tumor types listed above in Tumor Mutation Data, so *Ti* ∩ *Tj* = ∅; and for gene *g* ∈ *G* where *=* all sequenced genes \ O, where

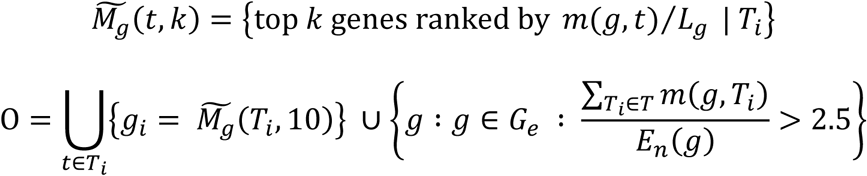

Compute the expected number of missense mutations in *g* in *Ti*, or

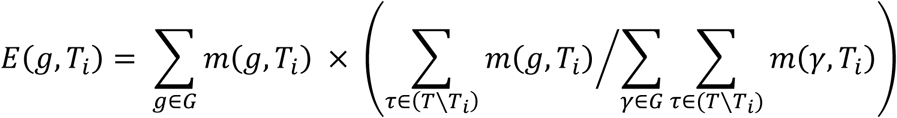

The calculated expected number of mutations for each gene was used to identify those genes under negative selection in one tumor type relative to the others (*N*). Genes were called asunder negative selection if they passed a fold-change and P value (calculated with the Poisson distribution) cutoff:

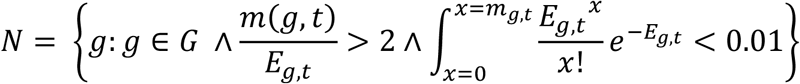

### Identifying pathways enriched in genes under purifying selection in specific tumor types

Pathways under purifying selection were identified from the list of genes under purifying selection generated after filtering out recurrent mutations as detailed above. The overlap between genes under purifying selection and a database of pathway gene sets (NDEx) (Pratt et al. 2015) was evaluated with a CDF of the hypergeometric distribution.

### Identifying gene sets under purifying selection in specific tumor types

Gene sets under purifying selection in specific tumor types were identified the same way as those gene sets under purifying selection in multiple tumor types (as detailed above), with the following differences. First, the expected number of mutations in each gene was estimated based on comparing one tumor type to other tumor types, as in “identifying genes under tumor type-specific selection,” above, where

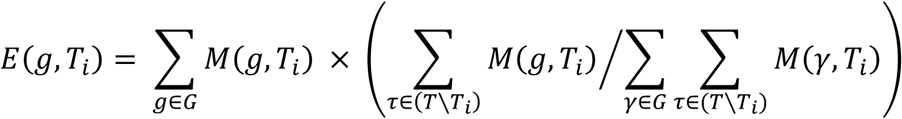

*T* = {melanoma, lung adenocarcinoma, and colorectal adenocarcinoma}

All other analysis of the depletion of mutations, gene set filtering, statistical and multiple hypothesis control was identical to “Identifying gene sets under purifying selection in multiple tumor types,” above.

For melanomas, gene sets were called to be under purifying selection if they had a q-value <0.05 and an observed / expected mutation ratio < 0.5.

For lung adenocarcinomas, gene sets were called to be under purifying selection if they had a q-value < 0.1 and an observed / expected mutation ratio < 0.55.

### Estimating the impact of sequencing more tumors

To evaluate the number of additional hits (individual genes identified as under purifying selection) we might find with more sequenced tumors, we down-sampled mutations by steps equivalent to the mutations of 40 average tumors in each tumor type, with 1000 replicates per down-sampling. The sampling was started from the dropping mutations equal to 80 random tumors and continued until the first step before the average number of hits returned was ≤ 1. Down-sampled data were fit to a four-parameter logistic curve (R^2^ ≥ 0.99):

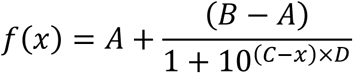

These fits were used to predict the number of new hits that would be found by steps of 10 additional sequenced tumors, and used to find an optimal distribution of sequenced tumors across tumor types to maximize the number of new hits.

### Determining the fraction of essential genes under purifying selection

Genes under purifying selection across tumor types (*Gp*) were defined as above, the union of genes in sets under purifying selection. Genes under increased purifying selection in melanomas 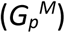 were defined similarly as the union of genes in sets under increased purifying selection in melanomas. Sets under increased purifying selection in melanomas were defined, above, in “Identifying gene sets under purifying selection in specific tumor types.” Gene sets were called to be under increased purifying selection in melanomas if they had a q-value < 0.05 and an observed / expected mutation ratio < 0.5.

Essential genes were identified from three CRISPR pooled screens (Hart et al. 2015; Wang et al. 2015; Tzelepis et al. 2016), as discussed in “Essentiality analysis of genes under purifying selection,” above. In each screen, a gene was called essential or not essential in each tested cell line. From this data, for each gene *g*, a score *Ci*(*g*) was recorded for screen *i*, or the number of tested cell lines in which this gene was deemed essential, based on the published results, in each screen. For each screen *i*, a gene was deemed essential if *Ci*(*g*) ≥ the number of cell lines tested in screen *i* - 2. The set of genes deemed essential in each screen *i* was then termed 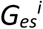. 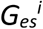 was also filtered so that it only included genes that were members of the sets in 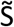 (the filtered gene sets from the Molecular Signature Database, see above), as those were the only genes that could be called under purifying selection.

The proportion of genes in 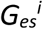 that were under selection (members of *Gp* or 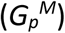 ) was then assessed.

### Code Availability

The code used in these analyses is available on request of the authors.

### Data access

All post-analysis data are included in this manuscript in supplemental tables. All data analyzed were obtained from other sources.

